# Repeated coastal observations and molecular confirmation of *Alatina grandis* (Cubozoa: Alatinidae) in the Gulf of Aqaba, Red Sea

**DOI:** 10.64898/2026.10.14.755470

**Authors:** Zafrir Kuplik, Ilka Straehler-Pohl, Anastasiia Iakovleva, Gil Koplovitz, Tamar Guy-Haim

## Abstract

Cubozoans are rarely documented in the Red Sea, and the taxonomic identity of large alatinid medusae previously reported from the Gulf of Aqaba has remained uncertain. Historical observations from Eilat in 1997 and 2000 were reported as *Carybdea alata*, although their exceptionally large size was recognized as being within the range of *C. grandis* (currently *Alatina grandis*); unresolved carybdeid taxonomy at the time prevented a definitive species-level assignment. Following more than two decades without documented observations, five large alatinid medusae were recorded on separate occasions along the Eilat coastline between July 2025 and August 2026. One specimen was collected, while high-resolution in situ photographs and video recordings of live medusae provided material for morphological assessment. Diagnostic morphological characters were consistent with *Alatina grandis*, and molecular analysis of the collected specimen independently confirmed its identity as *A. grandis*. These findings provide the first molecular confirmation of *A. grandis* from the Gulf of Aqaba and the Red Sea and establish the identity of the alatinid medusae documented during the recent series of observations. The clustering of five observations within one year, following more than two decades without documented occurrences, raises questions regarding the temporal occurrence and distribution of this rarely reported cubozoan in the northern Red Sea.

## Introduction

Cubozoans (Cnidaria: Cubozoa), commonly known as box jellyfish, comprise a relatively small class of gelatinous zooplankton. Nevertheless, they have attracted considerable attention because of their potent venom, sophisticated visual system, and active predatory behaviour (Bentlage et al., 2010; Kingsford & Mooney, 2014; Nilsson DE et al., 2005; Petie et al., 2011). However, despite their public notoriety and increasing scientific attention, the diversity, distribution, and biogeography of many cubozoan species remain incompletely understood, particularly in regions where records remain sparse. This knowledge gap is particularly evident in the Red Sea. Early accounts of Red Sea cubozoans were provided by Stiasny (1939), who reported carybdeid medusae from the region based on morphological observations. More than six decades later, Zakai and Galil (2001) documented two exceptionally large carybdeid medusae from the Gulf of Elat (Gulf of Aqaba), one observed alive in 1997 and another collected in 2000. The authors noted that the specimens were “clearly within the size range of *Carybdea grandis*” (currently *Alatina grandis*) but refrained from assigning them to species because the taxonomy of the *Carybdea alata* complex remained unresolved at the time. Consequently, the identity of these large alatinid cubozoans has remained uncertain for more than two decades.

The taxonomy of the family Alatinidae has undergone substantial revision during the past two decades. Gershwin (2005) established the genus *Alatina* and redefined the family Alatinidae. Subsequently, Straehler-Pohl and Gul (2017) revised the morphology and taxonomy of the family, providing updated diagnoses and identification characters for its constituent species. In parallel, integrative morphological and molecular studies demonstrated that several nominal species previously assigned to the *Carybdea alata* complex represent a single circumtropical species, *Alatina alata*, while confirming that *A. grandis* is a distinct species, genetically well separated from the *A. alata* complex (Bentlage et al., 2010; Lawley et al., 2016). Nevertheless, confirmed records of *A. grandis* remain uncommon, and its geographic distribution throughout the Indo-Pacific has not yet been fully resolved.

Following decades of only sporadic observations of large alatinid medusae in the Gulf of Aqaba, including a photographic record from 2014 that lacked sufficient morphological detail for species-level identification, five independent sightings were recorded along the Eilat coastline between July 2025 and September 2026 (Fig. 1; Table 1). This increase in the frequency of observations is particularly noteworthy given the extensive scientific activity, recreational diving, and tourism along the Eilat coastline.

**Table 1.**
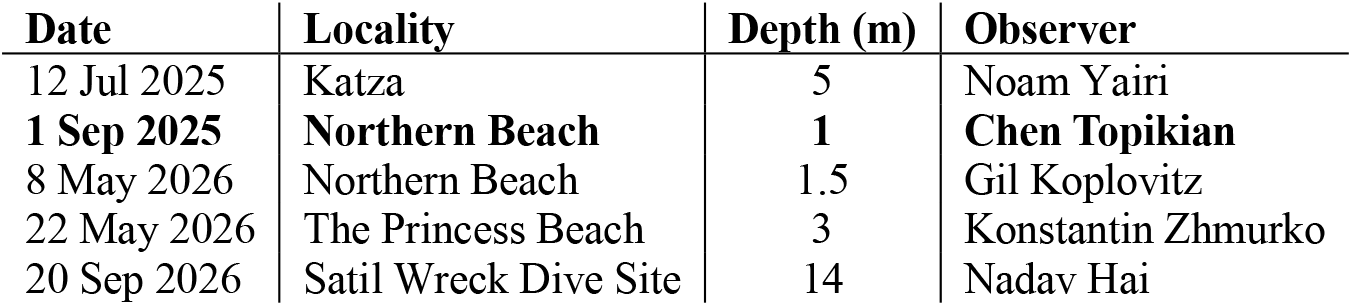
Details of the observed *A. grandis* along the coastal waters of Eilat, Gulf of Aqaba. Bold text indicates details for the specific specimen that was collected and used in the molecular analysis.

**Fig 1.**
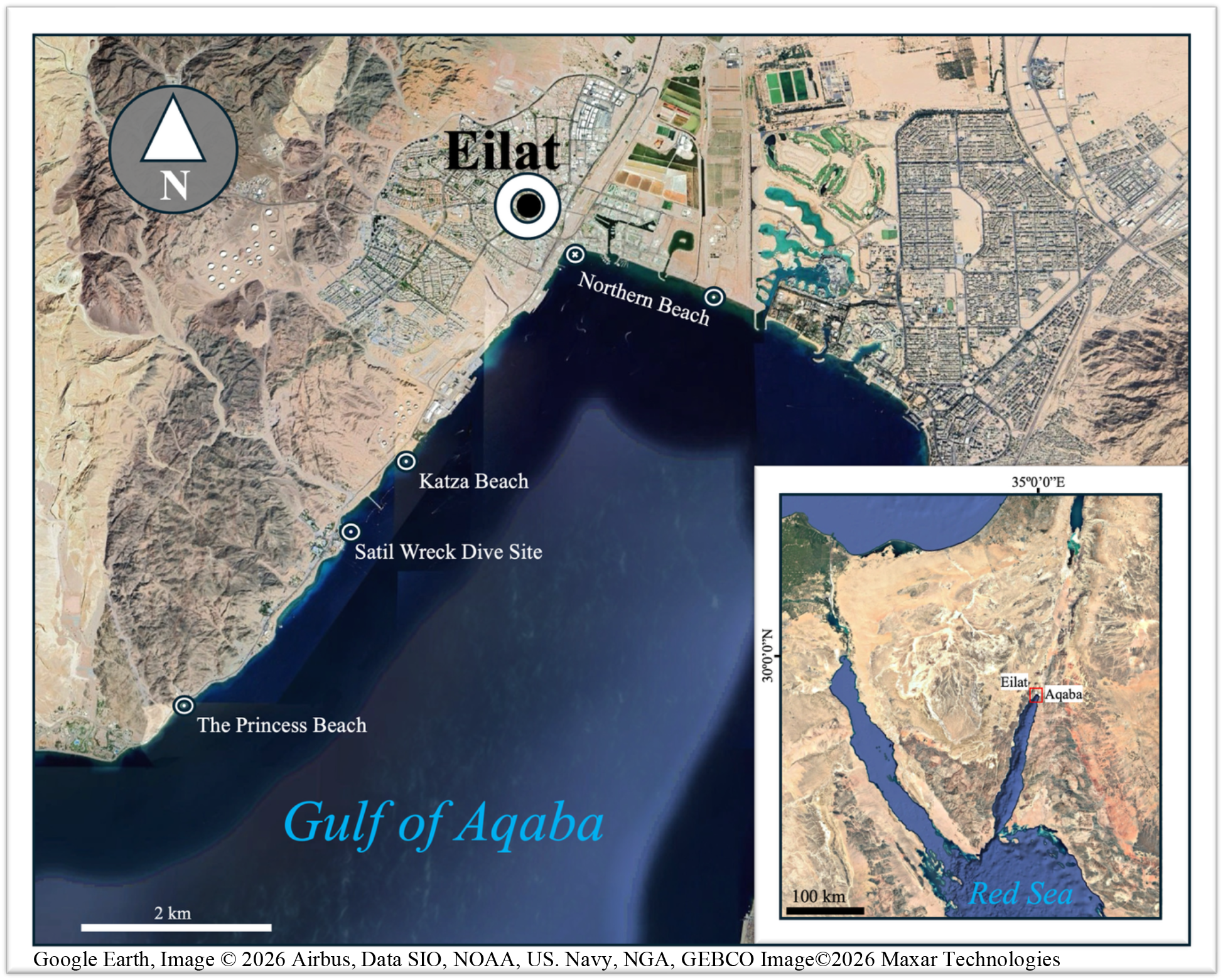
Map showing the sites of recent *Alatina grandis* observations along the Eilat coastline, Gulf of Aqaba, northern Red Sea. The circle in the Northern Beach containing an X indicates the collection site of the specimen used for molecular analysis, whereas the remaining circles represent observations documented by photographs or video recordings.

In the present study, we investigated these observations using an integrative taxonomic approach, combining morphological evidence from the collected specimen and high-resolution *in situ* photographs and video recordings of live medusae with molecular analyses of the collected specimen. Together, these complementary lines of evidence confirm the identity of the recently observed medusae as *A. grandis* and provide the first molecular confirmation of the species from the Gulf of Aqaba and the Red Sea.

## Materials and methods

### Specimens Examined

On 1 September 2025, a large alatinid medusa was collected from the wave-breaking zone along the northern shore of the Gulf of Eilat (Gulf of Aqaba, northern Red Sea), opposite Lifeguard Station No. 1, by Chen Topikian of the Israel Nature and Parks Authority. As no other preservation method was available immediately following collection, the specimen was initially stored frozen for subsequent morphological and molecular analyses. After thawing, the specimen was photographed, and bell and tentacle dimensions were measured using a ruler. The rhopalial niche ostium was also examined and photographed at this stage. The specimen was then fixed in 4% formalin and deposited in the Coelenterata Collection of the Steinhardt Museum of Natural History, Tel Aviv University, Tel Aviv (SMNHTAU-Co.39445). Although freezing and thawing caused substantial distortion and deterioration of the gelatinous tissues, several diagnostic morphological features remained sufficiently preserved for examination and documentation. Detailed inspection of the deposited specimen enabled the velarial canals to be photographed and characterized. All photographs of the collected specimen, including those of the whole specimen, rhopalial niche, and velarial canals, were taken using the camera of a Samsung Galaxy S22 smartphone.

An additional large alatinid medusa was photographed *in situ* by Gil Koplovitz of the Interuniversity Institute for Marine Sciences in Eilat (IUI) on 8 May 2026, along the northern shore of the Gulf of Eilat. Still images were obtained using a Sony A7RIII camera with a Sony 90 mm f/2.8 macro lens, Salted Line housing, Sea&Sea YS-D3 Duo strobe, and Nauticam MFO-3. The high-resolution images documented the live medusa in excellent condition and provided clear views of diagnostic morphological characters that were obscured or altered in the frozen specimen. Video footage of another live medusa, recorded by Konstantin Zhmurko on 22 May 2026 at approximately 3 m depth off Princess Beach, in the southernmost part of Eilat near the Israeli Egyptian border, was also examined to document morphological features visible *in situ*, including natural tentacle coloration. The bell length was estimated by the photographer to be approximately 25–30 cm and is provided as Supplementary Video S1. Taxonomic identification was based on morphological evidence from the collected specimen and the *in situ* photographic and video material, with molecular analyses of the collected specimen providing independent confirmation of the species identity.

### Molecular Analysis

Total genomic DNA was extracted from a dislodged tentacle using the InviSorb Spin Tissue Mini Kit (Invitek Diagnostics, Germany) according to the manufacturer’s specifications. Following the DNA extraction, the cytochrome *c* oxidase subunit I (COI) gene was amplified using PCR with universal primers LCO1490 and HCO2198 (Folmer, Black, Hoeh, Lutz, & Vrijenhoek, 1994) and 16S ribosomal DNA large subunit was amplified using the primers 16-L (Ender & Schierwater, 2003) and Aa_H16S_1541H (Bayha & Dawson, 2010). Reaction conditions for COI amplification were: 94 °C for 2 min, followed by 5 cycles of 94 °C for 40 s, 45 °C for 40 s, and 72 °C for 1 min, and followed by 30 cycles of 94 °C for 40 s, 51°C for 40 s, and 72 °C for 1 min, and a final elongation step of 72 °C for 10 min. Reaction conditions for 16S rRNA amplification were as follows: 95°C for 5 min followed by 35 cycles of 95°C for 1 min, 45°C for 1 min, and 72°C for 1 min, and a final elongation step of 72°C for 10 min. Obtained PCR products were separated on 1.5% agarose gel and stained with GelRed (Biotium Inc., USA). Purification and Sanger sequencing were done by Hy Laboratories Ltd (Rehovot, Israel). For each gene marker, forward and reverse sequences were assembled into a consensus contig using BioEdit 7.0.9 (Hall, 2004).

A total of 36 COI and 33 16S sequences of cubozoans were analyzed, including sequences of the *A. grandis* individual obtained in this study and *Alatina* cf. *grandis* from Saipan, Northern Mariana Islands in the western Pacific Ocean (Lawley et al., 2016). Sequence alignments were conducted using ClustalW embedded in MEGA v11.0 (Tamura, Stecher, & Kumar, 2021). The best-fitting substitution models were selected according to the Bayesian Information Criterion using Maximum-likelihood (ML) model selection in MEGA. ML analyses were performed using the T92+G+I model for COI sequences and GTR+G+I model for 16S sequences with 1000 bootstrapping replicates each.

## Results

SYSTEMATICS

Phylum CNIDARIA Hatschek, 1888

Class CUBOZOA Werner, 1973

Order CARYBDEIDA Gegenbaur, 1857

Family ALATINIDAE Gershwin, 2005

Genus *Alatina* Gershwin, 2005

*Alatina grandis* (Agassiz & Mayer, 1902)

### Morphology

The collected specimen had an approximate bell height of 21 cm. The total length of the preserved and contracted tentacle was estimated at approximately 28 cm, based on the combined length of the detached portion and the portion remaining attached to the pedalium (Fig. 2a, b), whereas the tentacles of the swimming medusa appeared considerably longer and were estimated by the photographer to exceed 1 m in length. A single pedalium was present at each of the four interradial corners of the bell, each bearing a single tentacle (Fig. 3a), an arrangement consistent with the diagnostic morphology of Carybdeida (Bentlage & Lewis, 2012). This arrangement was also clearly visible in the in-situ images of the live medusa (Fig. 3b). The rhopalial niche ostium was distinctly T-shaped (Fig. 4), a characteristic feature of Alatinidae (Gershwin, 2005; Bentlage & Lewis, 2012). Statoliths were not visible in the examined specimen, possibly owing to damage to the statocysts during freezing and thawing

**Fig 2.**
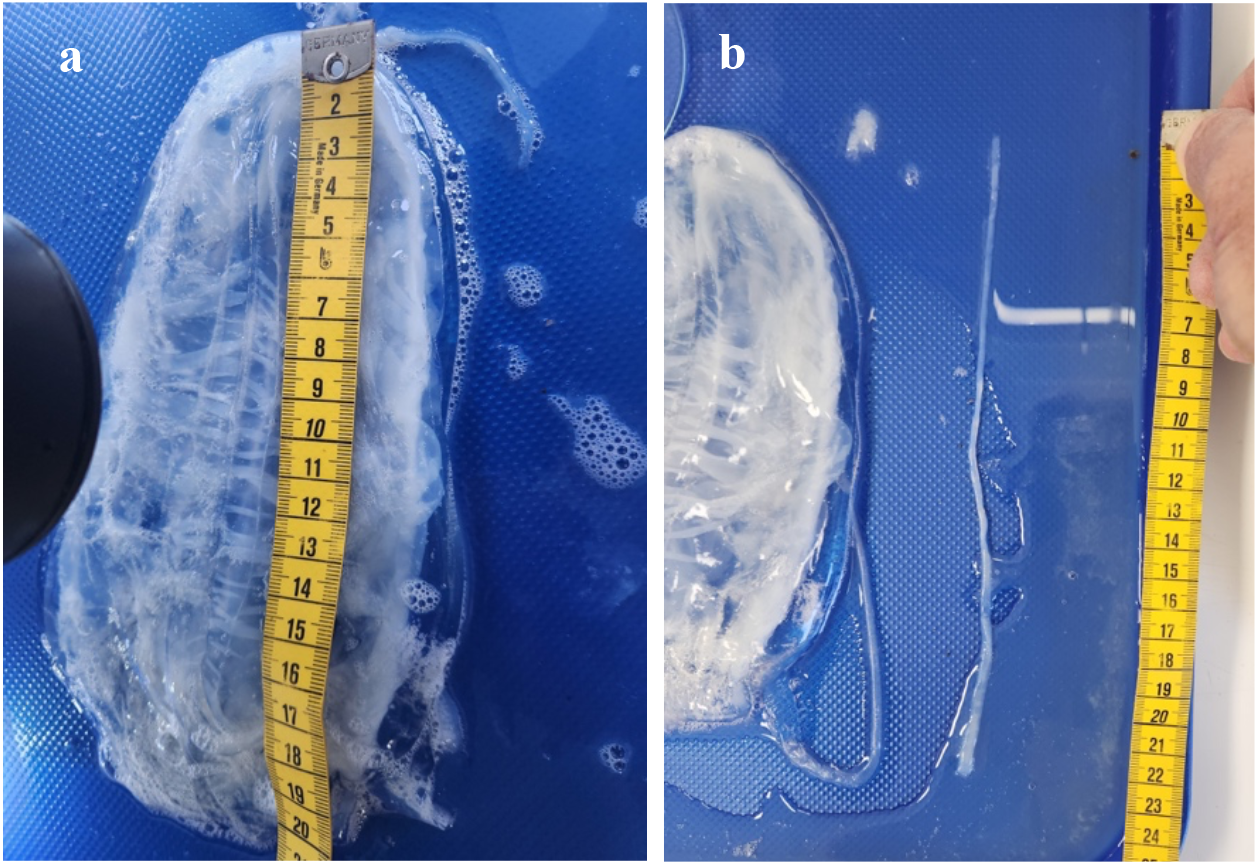
Measurements of the frozen *Alatina grandis* specimen collected from the northern Gulf of Aqaba. **(a)** Measurement of bell length. **(b)** Measurement of the detached tentacle and the tentacle portion attached to the pedalium, used together to estimate total tentacle length.

**Fig 3.**
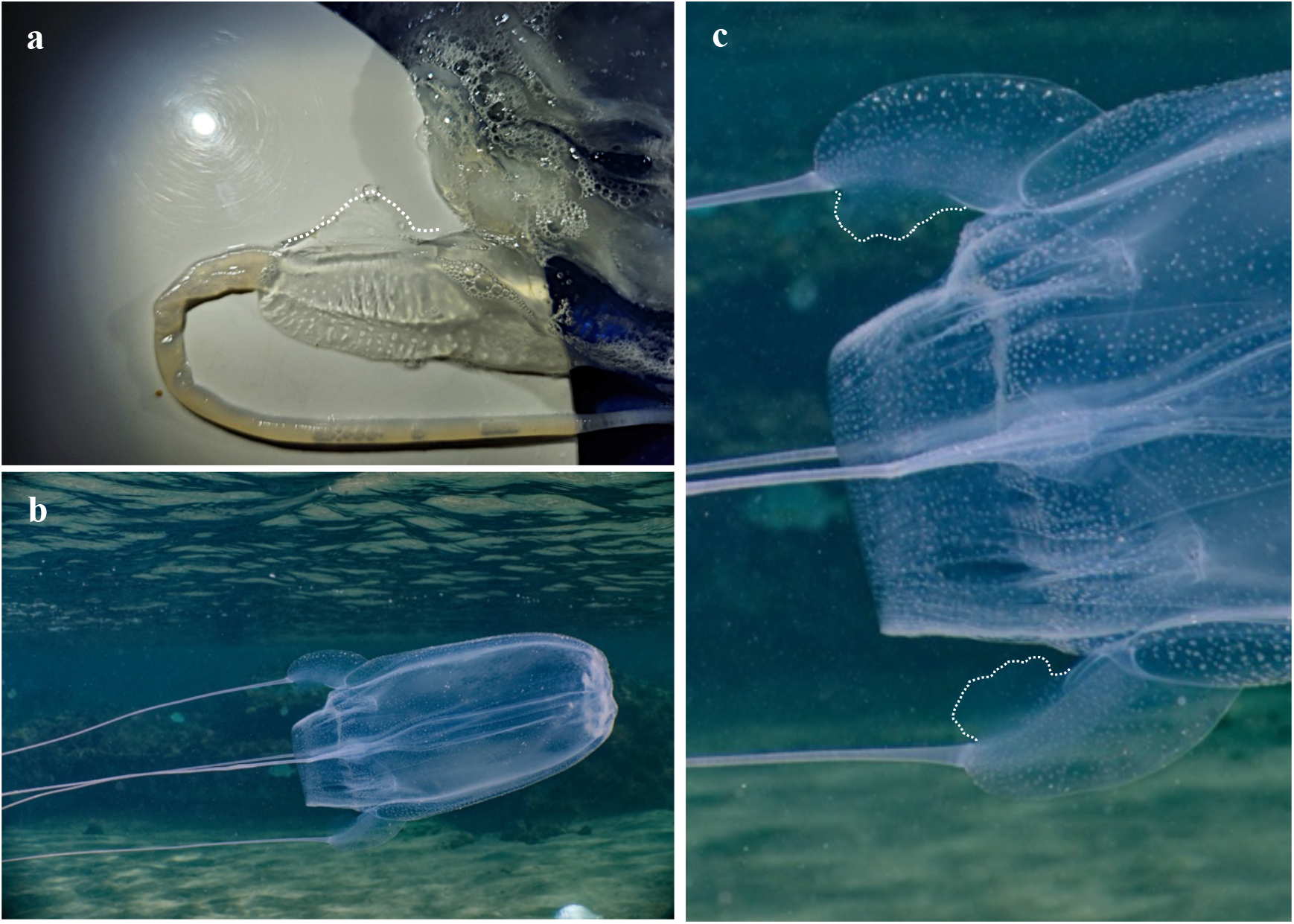
Pedalial morphology of *Alatina grandis*. (a) Pedalium of the collected specimen following thawing, showing the broad inner and outer pedalial wings and the single tentacle. (b) In situ photograph of the live medusa showing the four pedalia, each bearing a single tentacle. (c) Enlarged view of the pedalia of the live medusa (b), showing the broad, double-winged morphology; dotted lines indicate the margins of the inner pedalial wings. Photographs in (b, c) by Gil Koplovitz.

**Fig 4.**
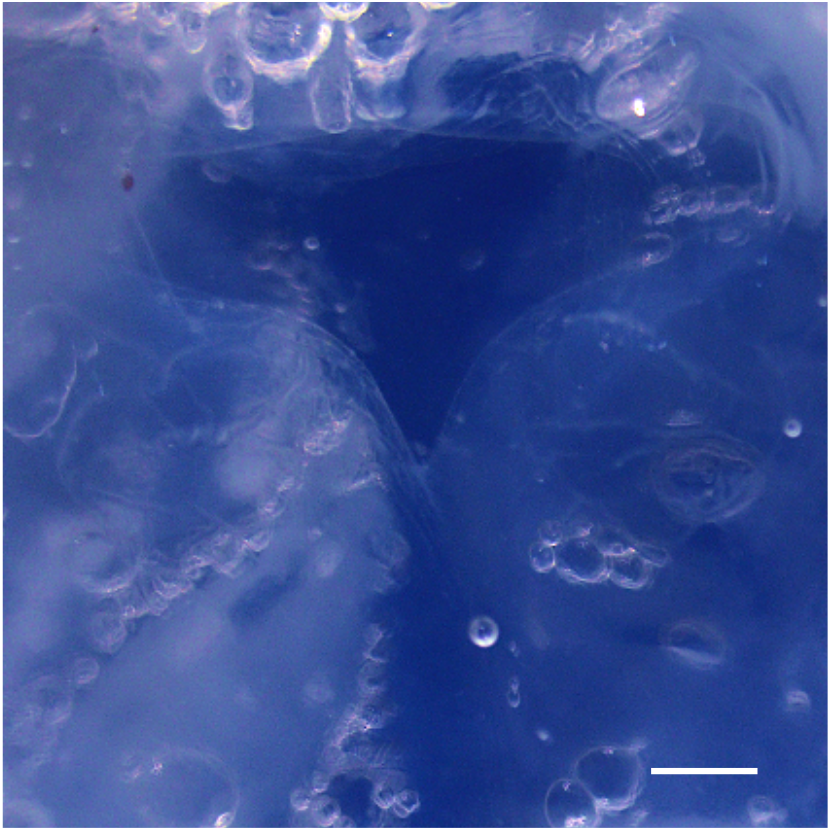
Rhopalial niche of the collected *Alatina grandis* specimen following thawing, showing the characteristic T-shaped rhopalial niche ostium. Statoliths are not visible. (Scale bar = 1000 µm)

At the species level, the pedalia were broad and double-winged, with the inner and outer wings of approximately equal width. This morphology was evident in both the collected specimen (Fig. 3a) and the in-situ images of the live medusa (Fig. 3b, c), with the margins of the inner pedalial wings highlighted in Fig. 3a and 3c.

The velarium exhibited three velarial canal roots per octant (Fig. 5). The canals arising from these roots were branched and bore numerous short, spike-like lateral projections and sharply pointed lobes, producing a distinctly jagged appearance. This morphology closely corresponds to the velarial canal pattern described for *Alatina grandis* (Straehler-Pohl & Gul, 2017). Additionally, the tentacles of the medusa documented on 22 May 2026 (Supplementary Video

**Fig 5.**
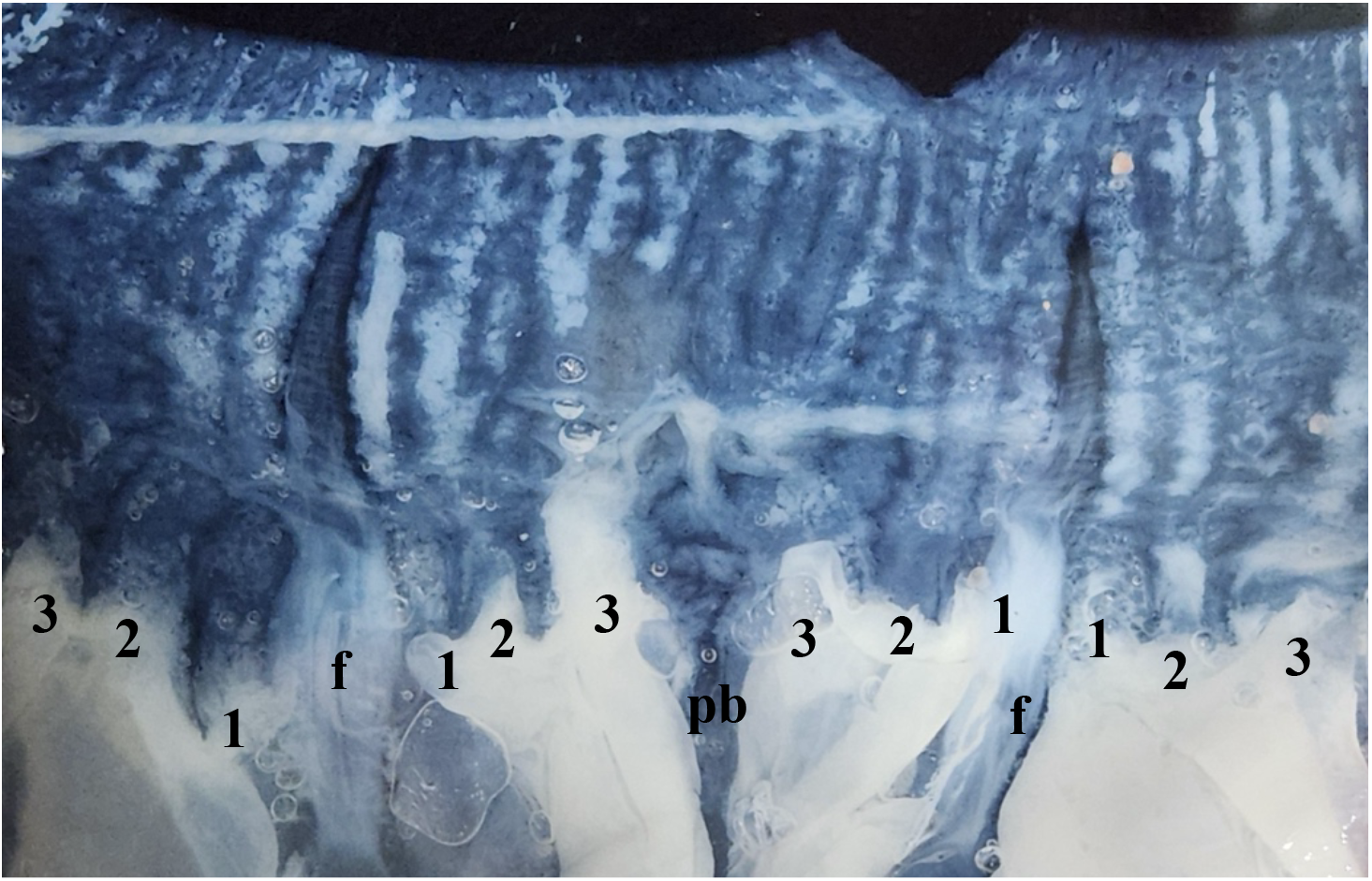
Velarial canal morphology of the collected *Alatina grandis* specimen following fixation in 4% formalin. Three adjacent octants are shown, delimited by the frenula (f), with a pedalium base (pb) centrally positioned. Each octant contains three velarial canal roots (1–3), giving rise to branched canals with short, sharply pointed lateral projections that produce a characteristic jagged appearance. The distance between adjacent frenula was approximately 5 cm in the fixed specimen.

S1) exhibited a distinct orange coloration, consistent with the coloration described for *A. grandis* by Gibbons et al. (2022).

### Molecular identification

The DNA barcode consisting of a fragment of 681 bp of the COI gene (PZ523033.1) and a 16S rRNA (PZ510948.1) fragment of 565 bp were sequenced from a medusa of *A. grandis*. The sequences were deposited in NCBI GenBank under the accession numbers (COI) and (16S). NCBI blastn yielded 80.11% and 84.66% identity to COI and 16S sequences of *A*. cf. *grandis* from Saipan. Maximum likelihood analysis of hydroid sequences obtained from GenBank (Figs. 6, 7) confirmed the identity of the Red Sea *A. grandis*.

**Fig 6.**
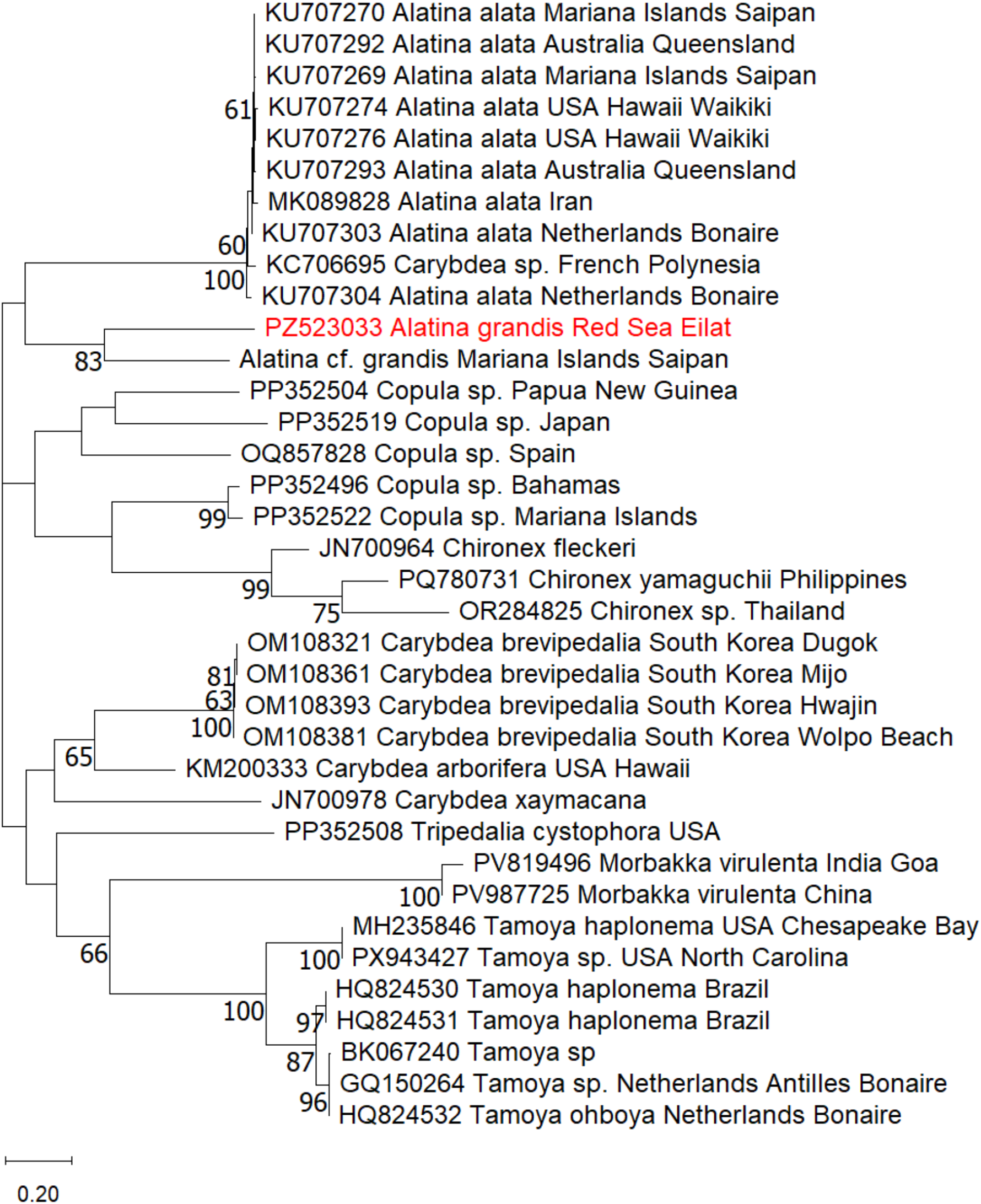
Maximum-Likelihood phylogenetic tree of *Alatina grandis* based on the COI gene, using the T92+G+I substitution model. The numbers below the branches indicate the percentage of ML bootstrap support (1000 replicates) for nodes that received at least 60% support. The scale bar denotes the estimated number of nucleotide substitutions per site.

**Fig 7.**
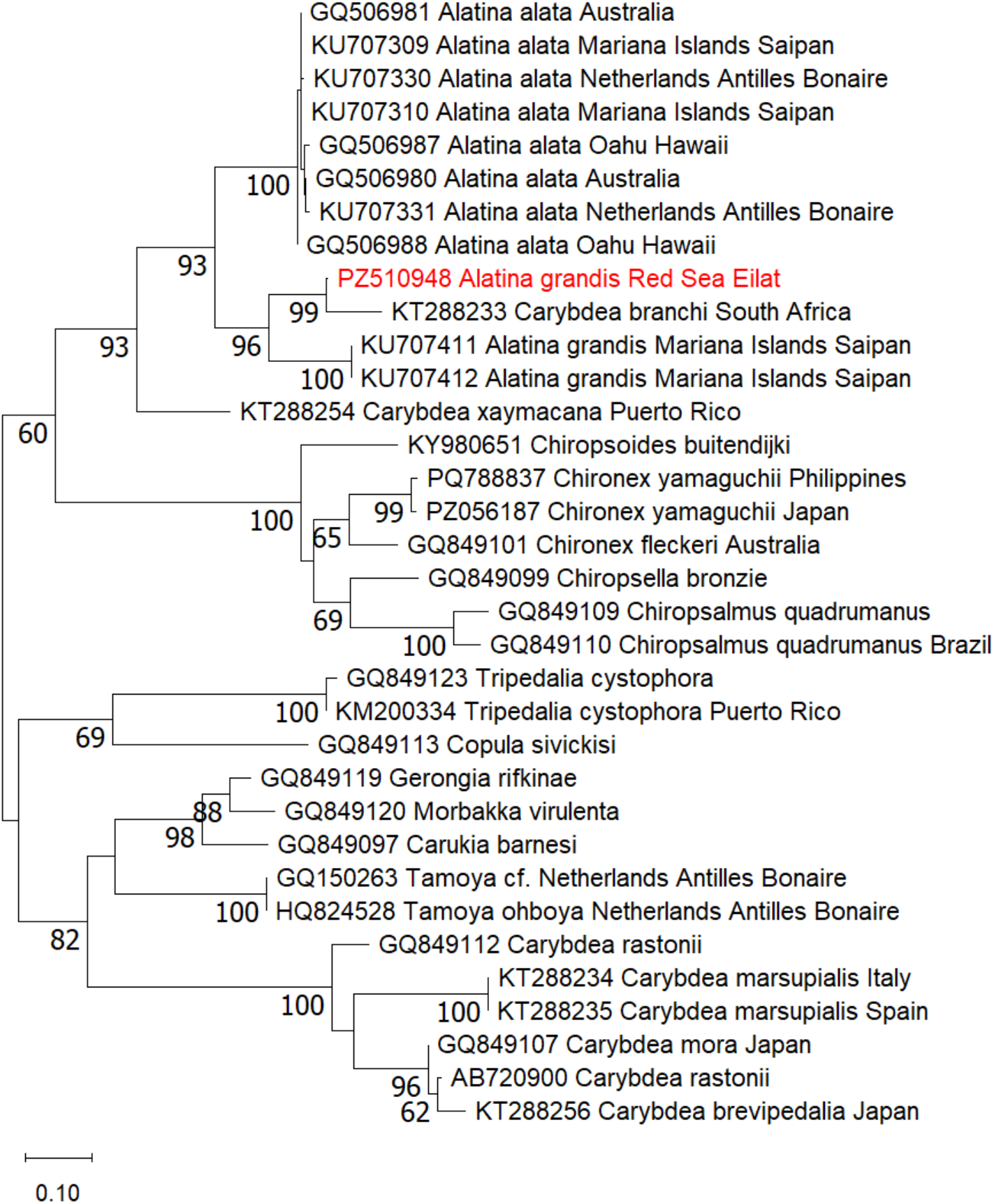
Maximum-Likelihood phylogenetic tree of *Alatina grandis* based on the 16S rRNA gene, using the GTR+G+I substitution model. The numbers below the branches indicate the percentage of ML bootstrap support (1000 replicates) for nodes that received at least 60% support. The scale bar denotes the estimated number of nucleotide substitutions per site.

## Discussion

The present study confirms the occurrence of *A. grandis* in the Gulf of Aqaba and provides the first molecular verification of the species from the Red Sea. Molecular identification of the collected specimen was supported by a suite of morphological characters documented both from the specimen itself and through *in situ* observations, including the T-shaped rhopalial niche ostium, broad double-winged pedalia, three velarial canal roots per octant with the characteristic complex, jagged branching pattern, and the distinct orange coloration of the tentacles reported for *A. grandis* by Gibbons et al. (2022). Importantly, morphological features consistent with *A. grandis* were documented across multiple observations rather than being restricted to the molecularly analyzed specimen. The recurrence of species-specific or characteristic features among separately observed medusae strengthens the interpretation that the recent records represent repeated occurrences of *A. grandis*, rather than isolated observations of unidentified alatinid medusae.

The occurrence of *A. grandis* in the Gulf of Aqaba is particularly noteworthy because the species is rarely documented throughout its known range. Its rediscovery in Pakistani waters followed a prolonged absence from the scientific record: a specimen collected near Karachi in 2014 was followed by six additional specimens, allowing the species to be redescribed and historical material from the John Murray Expedition of 1933–1934 to be recognized as *A. grandis* (Straehler-Pohl & Gul, 2017). A recent review of Cubozoa in the northern Indian Ocean (Jena and Sahu 2026) likewise lists *A. grandis* only from Pakistan and emphasizes the spatially and temporally fragmented nature of cubozoan records in the region. Thus, the present records extend the confirmed occurrence of this poorly documented species into the northern Red Sea and provide an important addition to its known distribution.

More intriguing than the confirmation of *A. grandis* itself is the recent increase in the frequency of observations along the Eilat coastline. The five recent sightings were distributed across approximately 14 months, rather than being confined to a single short-lived event. Together with the historical records, including a photographic observation from 2014 that could not be identified to species level, this pattern raises the possibility that large alatinid medusae have occurred intermittently in the Gulf for decades. An increase in documented records does not necessarily indicate population growth, particularly for cubozoans, for which long-term species-specific abundance data remain scarce (Jena & Sahu, 2026). Nevertheless, increased observation effort alone appears insufficient to explain the recent pattern in Eilat, where the coastline has been intensively observed through scientific activity, recreational diving, tourism, and underwater photography for decades. These observations may therefore reflect a genuine increase in the local abundance of *A. grandis*, a change in its spatial distribution that increased coastal encounters, or a combination of both processes.

Several biological and environmental mechanisms could account for this increased frequency of observations. One explanation that appears less likely is reproductive aggregation. Synchronized shoreward reproductive migrations are well documented in the congener *A. alata* in Hawaii, where medusae predictably enter shallow coastal waters following the full moon to reproduce (Chiaverano et al., 2013; Yanagihara et al., 2022). Individuals participating in these aggregations are sexually mature or recently spent, supporting their interpretation as spawning migrations. In contrast, none of the *A. grandis* medusae documented in Eilat showed visibly developed gonads. Although the reproductive biology of *A. grandis* remains largely unknown and the absence of conspicuous gonads cannot exclude all reproductive activity, the repeated occurrence of individuals without developed gonads provides no support for a spawning aggregation analogous to that of *A. alata*. Reproduction therefore appears unlikely to explain the recent increase in the frequency of coastal observations.

A more plausible biological mechanism is variation in food availability and consequent medusa survival or growth. Long-term observations of Hawaiian *Alatina* provide particularly relevant evidence. In a 14-year analysis of *Alatina moseri* (accepted as *A. alata*), interannual variation in medusa abundance was positively associated with primary production and zooplankton biomass, leading the authors to propose bottom-up regulation in which changes in primary production propagate through secondary production and prey availability to affect medusa abundance. This is particularly informative because reproductive timing alone could not explain the substantial variation in the numbers of *Alatina* appearing at the shoreline (Chiaverano et al., 2013). Comparable relationships have been reported for other cubozoans. The abundance and distribution of *Carybdea marsupialis* in the western Mediterranean have been associated with productivity-related variables, including chlorophyll-a and nutrients, as well as temperature and hydrodynamic conditions (Canepa et al., 2017). Juvenile abundance has also been positively associated with the availability of zooplankton prey, specifically cladocerns (Bordehore et al., 2020). Increased availability of suitable prey in the Gulf of Aqaba could therefore have enhanced growth or survival of young *A. grandis*, resulting in more individuals reaching conspicuous sizes and increasing their probability of detection. This hypothesis is biologically plausible but remains untested: the natural diet of *A. grandis* is poorly known, and no prey or zooplankton data have yet been linked directly to the Eilat observations. Determining whether zooplankton abundance or community composition differed during 2025–2026 from preceding years would therefore be particularly informative.

As in other medusozoans, cubozoans have a metagenetic life cycle comprising a benthic polyp stage and a pelagic medusa stage, both of which are affected by environmental conditions (Purcell, 2005). In cubozoans, temperature, salinity, and food availability have been experimentally shown to affect polyp survival, asexual proliferation, and metamorphosis (Courtney et al., 2016; Courtney & Seymour, 2013), providing a mechanism through which an inconspicuous benthic population may produce strongly varying numbers of medusae over time. Similarly, the seasonal appearance of juvenile *Carybdea marsupialis* has been associated with increasing spring temperature (Bordehore et al., 2020; Canepa et al., 2017). Thus, a resident *A. grandis* polyp population could conceivably have produced an unusually successful cohort during 2025–2026. The absence of developed gonads among the observed medusae is compatible with such a scenario, although it does not provide evidence for it. Critically, *A. grandis* polyps have never been documented in the Gulf of Aqaba, their habitat is unknown, and the environmental cues governing metamorphosis in this species have not been characterized. Whether the recent observations represent unusually successful recruitment therefore remains an open question.

Environmental conditions during the beginning of the observation period provide another potentially relevant clue. The Gulf of Aqaba experienced pronounced positive sea-surface temperature anomalies during summer 2025 relative to the NOAA Coral Reef Watch long-term climatology. A regional SST anomaly of +2.0°C was recorded on 29 June 2025, increasing to +3.0°C by 30 July, when SST reached 31.1°C (NOAA Coral Reef Watch, 2025a,b). Positive SST anomalies persisted throughout the summer of 2025. These conditions coincided approximately with the beginning of the recent sequence of *A. grandis* observations. The temporal correspondence is intriguing, particularly in light of the documented effects of temperature on cubozoan life-history processes (Bordehore et al., 2020; Canepa et al., 2017; Courtney et al., 2016; Courtney & Seymour, 2013) but should not be interpreted as evidence of causation.

In addition to the biological responses discussed above, anomalous thermal conditions may influence water-column stratification and circulation in the Gulf of Aqaba (Biton & Gildor, 2011; Carlson et al., 2014). This possibility is particularly relevant when considering the unusual habitat distribution reported for *Alatina*. Unlike most cubozoans, which are predominantly associated with coastal waters (Kingsford & Mooney, 2014), *Alatina* medusae have also been documented in offshore and deep-water environments (Lawley et al., 2016; Lewis et al., 2013), with *A. alata* recorded at depths of several hundred meters, including direct observations at approximately 540 m and collections from depths exceeding 1,000 m (Lewis et al., 2013; Lawley et al., 2016). Although comparable information on the vertical distribution of *A. grandis* is lacking, these observations raise the possibility that it too may spend substantial periods offshore or at depth and only occasionally occur within the shallow coastal zone. Notably, the Pakistani specimen central to the rediscovery of *A. grandis* was itself collected offshore by fishing trawl (Straehler-Pohl & Gul, 2017), providing at least some species-specific support for occurrence away from the immediate coastline. Such a distribution could make coastal occurrence particularly sensitive to physical transport. Wind and current direction, and current velocity have been associated with the abundance and spatial distribution of *C. marsupialis*, particularly among smaller medusae for which physical transport may be especially important (Canepa et al., 2017). Changes in circulation, convergence, stratification, or wind forcing in the northern Gulf could therefore conceivably transport *A. grandis* into the Eilat coastal zone or increase its retention there. Under this scenario, the recent observations would reflect an increase in coastal occurrence rather than population abundance. This could also reconcile the apparent paradox of a large and conspicuous medusa remaining virtually undocumented for decades along an intensively observed coastline: *A. grandis* may normally occur elsewhere in the Gulf, beyond the shallow coastal zone where observation effort is greatest. Observations of *A. alata* provide further support for an offshore-coastal connection in the genus. In Hawaii, *A. alata* is generally absent from shallow shoreline waters outside its predictable monthly appearances, while physical oceanographic processes have been implicated in its offshore distribution and subsequent shoreward occurrence (Yanagihara et al., 2022). These migrations differ fundamentally from the Eilat observations because they are associated with reproduction, whereas none of the *A. grandis* medusae documented here showed developed gonads. Nevertheless, they demonstrate that coastal occurrence may represent only a restricted component of the spatial ecology of *Alatina*. Thus, although reproductive migration appears unlikely to explain the Eilat observations, oceanographically mediated redistribution between offshore or deeper waters and the coastal zone remains a plausible hypothesis.

The mechanisms considered above are not mutually exclusive. Favorable environmental conditions could have enhanced medusa production, growth, or survival, resulting in a genuine increase in the local abundance of *A. grandis*, while changes in circulation may have increased the frequency with which medusae entered or remained within shallow coastal waters. The distribution of observations across multiple months and two consecutive years suggests that the recent records are unlikely to reflect a single, short-lived event. Rather, they may represent enhanced or repeated recruitment, improved medusa survival, recurrent transport into coastal waters, or a combination of these processes. Although the available evidence cannot distinguish among these possibilities, the historical records, including the putative observation from 2014, raise the possibility that *A. grandis* has persisted in the Gulf of Aqaba for decades. The recent increase in coastal encounters may therefore reflect changes in the abundance or spatial distribution of a long-established population rather than the arrival of a previously absent species. The available observations cannot presently distinguish among these mechanisms, but they identify several testable questions. Was zooplankton abundance or community composition unusual during 2025–2026? Did anomalous environmental conditions affect medusa production or survival? Does a resident benthic population of *A. grandis* occur in the Gulf, and if so, where? Do medusae normally occupy offshore or deeper waters? Were circulation or wind conditions during the observation period unusual in ways that favored transport or retention along the northern shoreline? Continued observations coupled with long-term environmental records, zooplankton sampling, current and wind measurements, and targeted searches for benthic stages could begin to distinguish increased recruitment or survival from physical redistribution.

At present, the recent records should therefore not be interpreted as evidence of a bloom, sustained population increase, or climate-driven range expansion. Rather, they document an unusual increase in the frequency of coastal encounters with a rarely recorded cubozoan. The absence of developed gonads argues against reproductive aggregation as an explanation, whereas food availability, environmentally mediated recruitment, and oceanographic redistribution remain plausible and potentially interacting mechanisms. Whether the recent observations represent an exceptional period or reveal a previously unrecognized aspect of the ecology of *A. grandis* remains unresolved. Continued monitoring of the Gulf of Aqaba, particularly beyond the intensively observed shallow coastline, will be essential for determining the distribution, seasonality, and environmental drivers of this poorly known species.

## Acknowledgements

This study was supported by ISF grant no. 1655/21 to T. Guy-Haim.

We thank Chen Topikian of the Israel Nature and Parks Authority for collecting and providing the *Alatina grandis* specimen used for molecular and morphological analyses, and Konstantin Zhmurko for documenting one of the medusae observed in Eilat and providing the video footage used in this study and as supplementary material. We are grateful to Allen Collins for sharing sequence data of Alatinidae.

## Data Availability Statement

The data underlying this article have been deposited in the GenBank Nucleotide Database at https://www.ncbi.nlm.nih.gov/genbank/, and are available from the corresponding author, [ZK], upon reasonable request.

## Competing interests

The authors declare no competing interests.

## Notes

### Competing Interest Statement

The authors have declared no competing interest.

### Summary of Updates

The revision is for a spelling fix: Koplovitz, instead of the Koplovich

